# Self-reassurance reduces neural and self-report reactivity to negative life events

**DOI:** 10.1101/2020.09.14.285486

**Authors:** Jeffrey J. Kim, Ross Cunnington, James N. Kirby

**Affiliations:** Compassionate Mind Research Group, School of Psychology, The University of Queensland, Brisbane, Queensland, Australia; The Centre of Advanced Imaging, The University of Queensland, Brisbane, Queensland, Australia

**Keywords:** Compassion, Self-Criticism, fMRI, Reassurance, Emotion

## Abstract

**Background:** Whilst research has shown how self-criticism may increase both neural and self-report markers of negative emotion, less well known is how self-reassurance - a compassionately-motivated cognitive self-relating style - may regulate negative emotion.

**Method:** Using fMRI, we invited participants to engage in self-criticism and self-reassurance toward written descriptions of negative life events (mistakes, setbacks, failures).

**Results:** Our results identify that neural markers of negative emotion and self-report markers of trial intensity during fMRI are suppressed under conditions of self-reassurance, relative to self-criticism.

**Limitations:** Future work to control for autobiographical memory during this fMRI task is needed, to explore how memory can contribute to self-reassurance and self-criticism.

**Conclusions:** Engagement in self-reassurance can reduce the ‘sting’ of negative life-events, both neural and self-report, which holds important implications for therapy.

## Introduction

Adverse life events are inescapable, be it a disruption in a career, dissolution of a relationship, or even a world-wide pandemic. These factors are known to take a toll on both physical and mental health outcomes (Solís et al., 2015) which can increase the likelihood of mortality (Puterman et al., 2020). These disappointments (e.g., making mistakes), losses (e.g., of hoped love) and fears (e.g., of rejection) are all triggers to self-criticism (Halamová et al., 2018; Kim et al., 2020). Indeed, self-criticism is a common self-relating style people use to cope, often resulting in an individual taking the frustration and anger out on themselves, which compounds the experience of pain psychologically and neurophysiologically (Kim et al., 2020). Whilst research has shown how self-criticism may increase both self-report (Cox et al., 2000; Gilbert et al., 2004) and neural (Longe et al., 2010; Lutz et al., 2016) markers of negative emotion, less well known is how self-reassurance - a compassionately-motivated cognitive self-relating style - may regulate how the brain responds toward negative life events.

Here we conducted an fMRI experiment which examined two distinct self-relating styles, self-criticism and self-reassurance (Kim et al., 2020; Petrocchi et al., 2019), when participants imagined themselves responding to mistakes, setbacks or failures. Importantly, we designed our experiment to deliberately tease apart neural markers of negative emotion, first by manipulating an emotional – neutral contrast at the first level of fMRI analysis, and explored how this activation may differ across self-criticism and self-reassurance. To anticipate our findings, we identified that how the brain encodes negative emotion differs under conditions of self-criticism versus self-reassurance, specifically showing how self-reassurance can down-regulate neural markers of negative emotion.

## Methods

Whilst the program of research within the present paper has been reported on previously, this examined the (neuro)physiological correlates of a brief, two-week compassion training paradigm (Kim et al., 2020). Here we focus on the novel whole brain markers of criticism and reassurance which have not been reported. As our fMRI method as reported in the previous paper is also the same imaging method used for the present paper, we have reproduced this section for clarity under a CC BY open access licence.

### Participants

40 participants (Mean age = 22 years, SD = .49, 27 female) took part in the present study. A University’s Ethics Sub-Committee approved the experimental protocol, and this project complies with the provisions contained in the *National Statement of Ethical Conduct in Human Research* and complies with the regulations governing experimentation on humans. Participants provided informed and voluntary, written and/or electronic consent.

### fMRI Stimuli

We created 60 written stimuli in total, consisting of a personal mistake, setback or failure. 30 statements were of emotional valence whereas 30 were neutral (i.e., “I fail to keep up with my commitments in life”, and “I keep up with my commitments in life”, respectively). Our neutral stimuli were created to describe a non-emotive, non-intense control to counterbalance the emotional stimuli set. For both emotional and neutral sets we assessed two metrics, valence (1-5, where 1 = Very Unpleasant) and intensity (1-5, where 1 = Not Intense). Our emotional statements (*n* = 30) were revealed to be sufficiently unpleasant (*M* = 1.89) and intense (*M* = 3.54), with all neutral statements (*n* = 30) described as less unpleasant (*M* = 3.80) and comparatively not intense (*M* = 2.34).

### fMRI Design

Within the scanner we examined participant’s neural responses to the validated (affective and neutral) written stimuli when engaged in self-criticism and self-reassurance (Figure 1). Participants were given one practice block with 16 trials (8 emotional and 8 neutral) repeated for self-criticism and self-reassurance, and made button press ratings on perceived trial intensity. After each trial within a block of either self-criticism or self-reassurance, participants rated how intense their degree of self-criticism or self-reassurance was to each statement (button-press on an MR-compatible button box which ranged from 1-4, where 1 = not very intense, and 4 = very intense). A typical trial consisted of stimuli presented for a 6 second duration, followed by a rating of intensity for a 3 second duration, and an inter-trial-interval of .5 seconds. The first order of instruction for a particular block, that is, self-reassurance verses self-criticism, was counterbalanced for a total of 8 blocks. As our focal contrast, we manipulated the emotionality of the statements within scan runs (“emotive” vs “neutral”), in a counterbalanced order across participants. 30 statements were quasi-randomized across participants and presented for a total of 30 trials per fMRI run (~6.5 min total duration) over a total of 8 repeated fMRI runs. Participants were given 10 practice trials of emotional and neutral stimuli, and rated stimuli on intensity.

**Figure 1.**
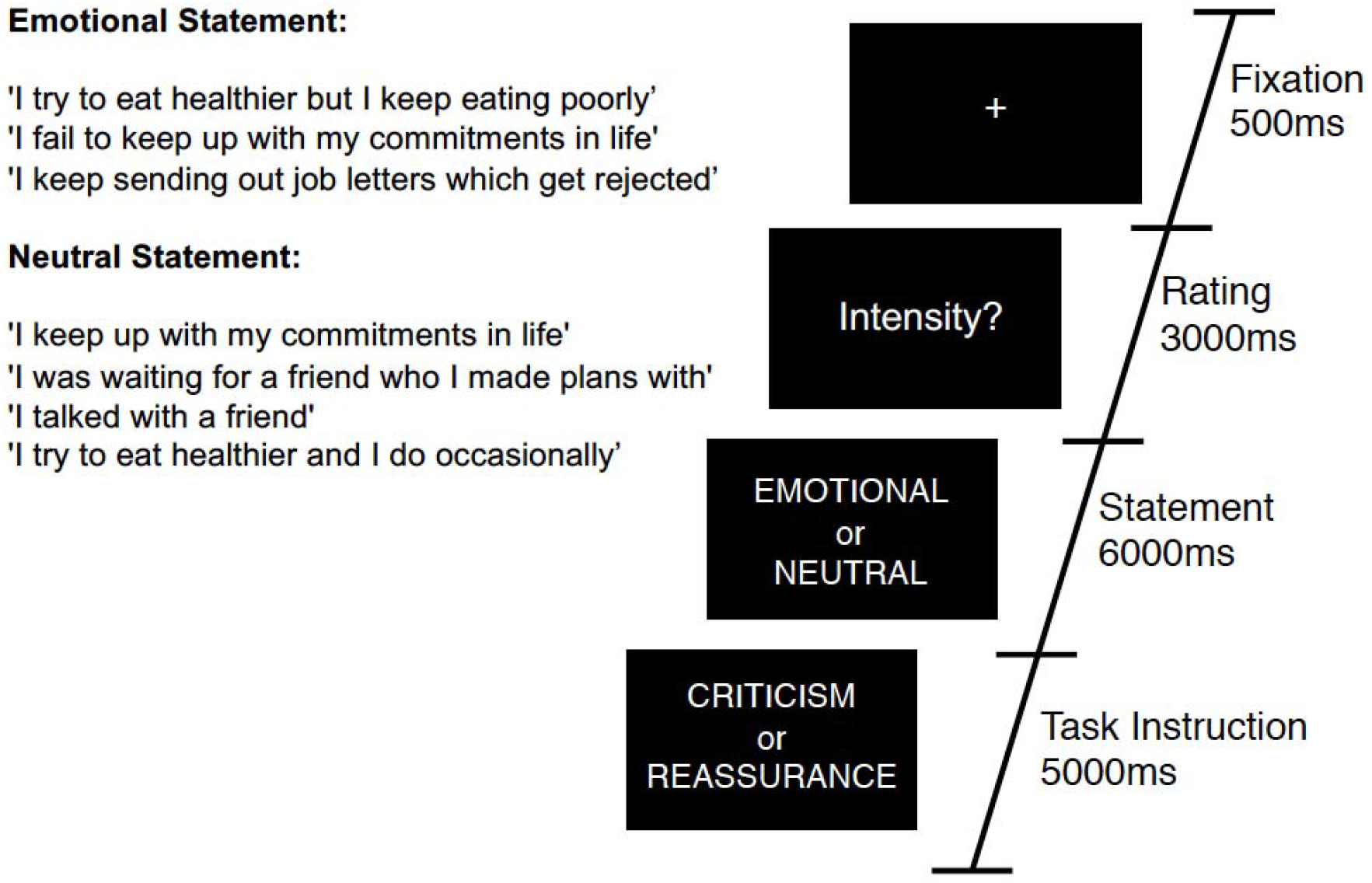
Task diagram for a typical trial. Participants were presented with 30 alternating trials of emotional or neutral statements which describe a mistake, setback or failure. Across 8 scan runs of 6 minutes each, participants were asked to engage with these statements from two different perspectives – four blocks of self-criticism, and four blocks of self-reassurance (order counterbalanced across participants). Example statements are presented inset.

### fMRI Acquisition and Pre-Processing

We collected our fMRI data on a Siemens Magnetom Prisma Fit 3 Tesla MRI scanner utilizing a 64-channel head-coil. A gradient-echo, echo-planar imaging sequence was used to acquire functional images, with the following sequence parameters: 60 horizontal slices (2 × 2-mm in-plane voxel resolution and 2-mm slice thickness plus 10% gap), repetition time (TR) 1000 ms; echo time (TE) 30 ms. Eight identical fMRI runs of 292 images (6 minutes each) were acquired. A 3D high-resolution, unified and denoised T1-weighted MP2RAGE image across the entire brain was also acquired and used as anatomical reference for subsequent pre-processing in SPM12 (TR = 4000 ms, TE = 2.93 ms, FA = 6°, 176 cube matrix, voxel size = 1-mm). Functional imaging data were pre-processed and analyzed using SPM12, implemented in MATLAB. Structural T1-scans were co-registered to the average of the spatially realigned functional slices. Next, an inbuilt segmentation routine was applied to register each structural T1-image to the standard MNI template in MNI space. These transform parameters elicited from segmentation were subsequently applied to all realigned images, resliced to a 2×2×2-mm resolution and smoothed with 6-mm full-width-at-half-maximum (FWHM) isotropic Gaussian kernel.

### fMRI First and Second-Level Analyses

For first-level data analysis, block-related neural responses to stimuli were modelled as 2 separate conditions (all combinations of emotional/neutral, self-criticism/self-reassurance) and convolved with the canonical hemodynamic response function (HRF). For group level analysis, whole-brain contrasts of self-criticism (emotional-neutral) stimuli were reported at a cluster-level threshold of *p* < .05, corrected for family-wise error, with clusters formed with a voxel-level height threshold at *p* < .001, uncorrected, with a cluster extent threshold of K = 144. Whole-brain contrasts of self-reassurance (emotional-neutral) stimuli were reported at a cluster-level threshold of *p* < .05, corrected for family-wise error, with clusters formed with a voxel-level height threshold at *p* < .001, uncorrected, with a cluster extent threshold of K = 144. Whole-brain repeated-measures contrasts of self-criticism (emotional-neutral) - self-reassurance (emotional-neutral) stimuli were reported at a cluster-level threshold of *p* < .05, corrected for family-wise error, with clusters formed with a voxel-level height threshold at *p* < .001, uncorrected, with a cluster extent threshold of K = 110. Brain regions shown to be significant had their anatomical labels identified with the Automated Anatomical Labelling (AAL) toolbox implemented in SPM12.

### Self-report markers of intensity during fMRI

Analysis of participant’s mean level of intensity ratings for reassuring trials (emotional stimuli: *M* = 2.45, *SD* = 0.48, neutral statements: *M* = 2.63, SD = 0.64) and critical trials (emotional stimuli: *M* = 2.92, *SD* = 0.45, neutral stimuli: *M* = 2.07, *SD* = 0.52) revealed intensity ratings were significantly higher for critical (emotional – neutral) but not for reassuring (emotional – neutral) trials (t(38) = 7.300, *p* < 0.001, and t(38) = −1.372, *p* = 0.178, *ns*, respectively).

## Results

First, group-level one-sample t-tests of the whole brain contrasts of emotional - neutral stimuli were conducted overall. We then examined how this contrast differed across self-criticism and self-reassurance. For neural markers of negative emotion during self-criticism, we observed activation in the “salience” (midcingulo-insular), “default-mode” (medial frontoparietal), and the occipital network (Uddin et al., 2019). Whilst neural markers of negative emotion during self-reassurance recruited activation in regions such as the medial-prefrontal cortex (MPFC) and visual cortex, we observed no activation of the salience network as shown under self-criticism. Across both these contrasts, clusters were formed at a cluster-level threshold of *p* < .05, corrected for family-wise error, with clusters formed with a voxel-level height threshold at *p* < .001, uncorrected (cluster extent threshold K = 144).

We next conducted a repeated-measures contrast between self-criticism (emotional – neutral) minus self-reassurance (emotional – neutral), as a marker of neural markers of negative emotion which differs between these two self-relating styles. Here, we identified brain activation across bilateral hippocampus (with a cluster which also included left putamen and left insula), thalamus, ACC, and occipital lobe, revealing neural markers of negative emotion are driven by self-criticism but not self-reassurance (cluster-level threshold of *p* < .05, corrected for family-wise error, with clusters formed with a voxel-level height threshold at *p* < .001, uncorrected, with a cluster extent threshold of K = 110). A repeated-measures contrast between self-reassurance (emotional – neutral) minus self-criticism (emotional – neutral) returned non-significant. Our experimental design is shown in Figure 1, and Figure 2 depicts the whole-brain results. Tables which present the thresholded output for each contrast are available online (https://osf.io/9wzu4/?view_only=91484c009daa4b03b676cbfa70940a6f).

**Figure 2.**
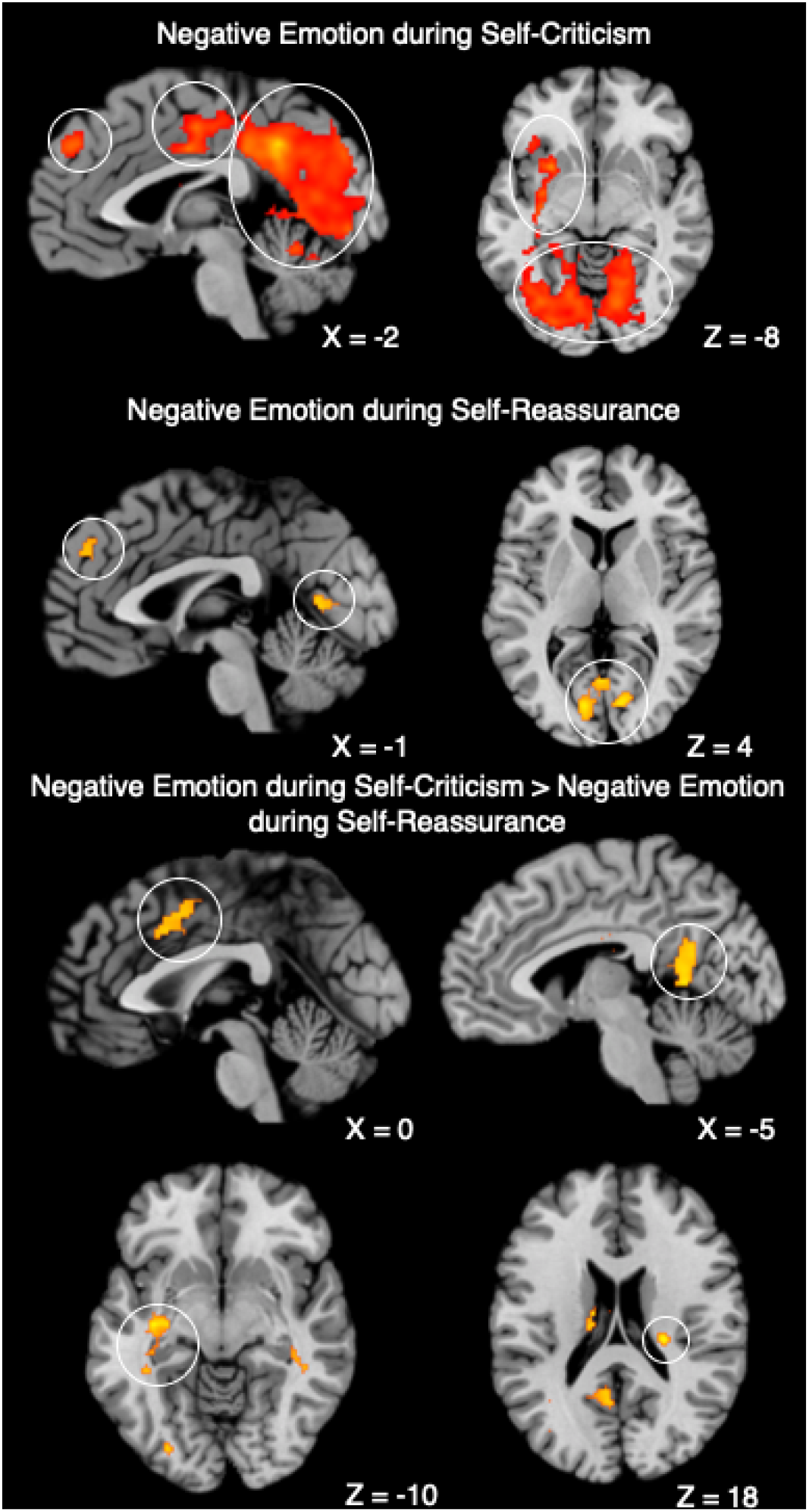
Neural markers of negative emotion across self-criticism and self-reassurance. **Negative Emotion during Self-Criticism: Left.** Sagittal image of MPFC (Left Circle), ACC (Middle Circle), and Left Lingual Gyrus and Cerebellum (Right Circle). **Right.**Axial image of Subcortical Regions (Top Circle) and Bilateral Visual Cortex (Bottom Circle). **Negative Emotion during Self-Reassurance: Left.** Sagittal image of MPFC (Left Circle) and Visual Cortex (Right Circle). **Right.** Axial image of Visual Cortex. **Negative Emotion during Self-Criticism – Negative Emotion during Self-Reassurance: Top Left.**Sagittal image of ACC. **Top Right.** Sagittal image of posterior cingulate. **Bottom Left.** Axial image of left putamen. **Bottom Right.** Axial image of Right Hippocampus. Coordinates reported in MNI-space. *N* = 40.

## Discussion

Here we investigated neural markers of negative emotion when participants engaged in self-criticism and self-reassurance toward negative life events (i.e., mistakes, setbacks or failures). Across both self-criticism and self-reassurance, our fMRI study revealed common activation across diverse regions such as the visual cortex (associated with mental imagery), salience network (associated with processing pain and threat), and default-mode network (associated with self-referential thought) (Uddin et al., 2019). Brain activation overall was more extensive for self-critical than self-reassuring trials, even though both contrasts did activate similar regions such as the MPFC and visual cortex. Furthermore, self-reassurance did not activate regions such as the insula, anterior cingulate cortex and amygdala. In addition, self-report ratings of intensity for emotional stimuli were supressed for self-reassuring versus self-critical trials. Importantly, a contrast of negative emotion between self-criticism minus self-reassurance revealed brain activation in regions such as the anterior cingulate cortex, insula and hippocampus. Taken together, our data show that neural and self-report markers of negative emotion are supressed during self-reassurance compared with self-criticism, providing evidence for how cultivating a reassuring self-relating style can regulate neural markers of negative emotion.

Whilst recruitment of the insula and anterior cingulate cortex have previously been shown for self-criticism (Hooley et al., 2012; Lutz et al., 2016), it is important to remark on bilateral hippocampus activation within the current experiment, which may be an indicator of autobiographical memory recall (Aly, 2020; McCormick et al., 2020). Given our paradigm instructions were for participants to engage in self-critical thoughts from the stimuli presented, it is entirely possible that for reference participants engaged in their own first-person accounts from situations in their own lives (Holland & Kensinger, 2010). However, given we did not explicitly control for autobiographical memory, future work is needed to explore how memory may contribute to neural markers of self-criticism or self-reassurance. To position our results in the broader literature on the neuroscience of empathy and compassion, we have shown that brain regions for processing negative emotion toward others (Ashar et al., 2017; Kim, Cunnington & Kirby, 2020; Zaki et al., 2016) were shown to not be recruited during compassion to the self. Specifically, we have shown that neural markers of negative emotion are suppressed during attempts to be compassionate and reassuring to one’s suffering. Our data suggest that engagement in self-reassurance is a way to reduce the ‘sting’ of negative life-events, both neural and self-report, which is a timely finding in our current global environment.

## Author Contributions

JJK, RC and JNK conceived and designed the experiment; JJK acquired, analysed and interpreted the data; JJK and JNK wrote the manuscript. All authors have given final approval for submission.

## Acknowledgements

JJK thanks Fernanda L. Ribeiro for helpful feedback and comments on a manuscript draft.

## Role of the Funding Source

This research was supported by a University of Queensland Research and Teaching fellowship awarded to JNK. JJK was supported by an Australian Postgraduate Scholarship.

## Competing Interests Statement

The authors declare no competing interests.

## Notes

### Competing Interest Statement

The authors have declared no competing interest.

