## Supplementary Tables for "Self-reassurance reduces neural and self-report reactivity to negative life events"

| Area | Cluster peak co-ordinates (XYZ) | Peak t-score | $K_E$ | $P_{FWE\text{-}corr}$ |
|---|---|---|---|---|
| *Reassurance: Emotional – Neutral* | | | | |
| Left Angular Gyrus | -40 -56 36 | 5.53 | 244 | .001 |
| Left Inferior Parietal | | | | |
| Left Lingual Gyrus | -10 -82 4 | 5.23 | 305 | .001 |
| Left Calcarine | | | | |
| Right Lingual | | | | |
| Right Lingual | 12 -80 4 | 5.07 | 299 | .001 |
| Right Calcarine | | | | |
| Right Cerebellum | | | | |
| Left Frontal Supp Medial | 2 50 34 | 4.44 | 147 | .018 |
| Right Frontal Supp Medial | | | | |

***Table 1.*** Brain regions that showed significantly greater activation for self-reassurance (emotion - neutral). Peak thresholded at FWE, $p$ <0.05 with K = 144 and coordinates reported in MNI-space. Cluster labelling was used from the AAL toolbox implemented in SPM12.

| Area | Cluster peak co-ordinates (XYZ) | Peak t-score | $K_E$ | $P_{FWE\text{-}corr}$ |
|---|---|---|---|---|
| *Criticism: Emotional – Neutral* | | | | |
| Left Calcarine | -4 -52 36 | 9.21 | 11295 | .001 |
| Right Lingual | | | | |
| Left Lingual | | | | |
| Left Precuneus | | | | |
| Right Calcarine | | | | |
| Left Fusiform | | | | |
| Left Cuneus | | | | |
| Left Middle Cingulate | | | | |
| Right Cuneus | | | | |
| Left Inferior Occipital | | | | |
| Right Fusiform | | | | |
| Right Precuneus | | | | |
| Right Posterior Cingulate | | | | |
| Left Cerebellum | | | | |
| Right Cerebellum | | | | |
| Vermis | | | | |
| Right Middle Cingulate | | | | |
| Right Posterior Cingulate | | | | |
| Left Inferior Temporal | | | | |
| Vermis | | | | |
| Left Crusl 1 Cerebellum | | | | |
| Left Supplementary Motor Area | | | | |

| | | | | |
|---|---|---|---|---|
| Left Parahippocampal | | | | |
| Left Hippocampus | | | | |
| Left paracentral Lobule | | | | |
| Right Inferior Occipital | | | | |
| Right ParaHippocampal | | | | |
| Right Cerebellum Crusl 1 | | | | |
| Left Middle Temporal | | | | |
| Right Cerebellum 3 | | | | |
| Right Middle Cingulate | | | | |
| Left Putamen | -28 19 -14 | 7.45 | 677 | .001 |
| Left Insula | | | | |
| Left Inferior Orbital | | | | |
| Left Frontal Inferior Tri | | | | |
| Left Hippocampus | | | | |
| Left OFCpost | | | | |
| Left Amygdala | | | | |
| Left Inferior Temporal | | | | |
| Right Caudate | 10 0 22 | 6.02 | 349 | .001 |
| Right Insula | | | | |
| Right Frontal Inferior Oper | | | | |
| Right Putamen | | | | |
| R Rolandic Operculum | | | | |
| Left Frontal Superior Medial | -4 46 36 | 5.72 | 145 | .026 |
| Right Frontal Superior Medial | | | | |
| Left Caudate | -8 -18 22 | 5.62 | 350 | .001 |
| Left Thalamus Putamen | | | | |
| Left Thalamus LP | | | | |
| Left Angular | -44 -46 22 | 5.45 | 494 | .001 |
| Left SupraMarginal | | | | |
| Left Mid Occipital | | | | |
| Left Mid Temporal | | | | |
| Left Sup Temporal | | | | |
| Left Inferior Parietal | | | | |
| Left Precentral | -30 -28 48 | 5.26 | 625 | .001 |
| Left Postcentral | | | | |
| Left Parietal Sup | | | | |
| Left Paracentral Lobule | | | | |

***Table 2.*** Brain regions that showed significantly greater activation for self-criticism (emotion - neutral). Peak thresholded at FWE, $p$ <0.05 with K = 144 and coordinates reported in MNI-space. Cluster labelling was used from the AAL toolbox implemented in SPM12.

| Area | Cluster peak co-ordinates (XYZ) | Peak t-score | $K_E$ | $P_{FWE-corr}$ |
|---|---|---|---|---|
| *Criticism (Emotional – Neutral) – Reassurance (Emotional – Neutral):* | | | | |
| Left Putamen | -36 -16 -10 | 5.33 | 314 | .001 |
| Left Hippocampus | | | | |
| Left Pallidum | | | | |
| Left Insula | | | | |
| Right Hippocampus | 24 -24 18 | 4.49 | 147 | .001 |
| Left Thalamus VL | -12 -12 18 | 4.46 | 158 | .001 |
| Left Caudate | | | | |
| Left Thal VPL | | | | |
| Left Thal LP | | | | |
| Left Precuneus | -4 -56 4 | 256 | 256 | .001 |
| Left Calcarine | | | | |
| Left Lingual | | | | |
| Vermis_4_5 | | | | |
| Left Posterior Cingulate | | | | |
| Left Cerebellum | | | | |
| Left Cuneus | | | | |
| Left Inferior Occipital | -28 -84 -4 | 117 | | .001 |
| Left Middle Occipital | | | | |
| Left Middle Cingulate | 0 14 36 | 115 | | .001 |
| Right Middle Cingulate | | | | |
| Left Supp Motor Area | | | | |
| R Supp Motor Area | | | | |
| Left Middle Occipital | -38 -72 24 | 127 | | .001 |
| Left Angular | | | | |
| Left Middle Temporal | | | | |

***Table 3.*** Brain regions that showed significantly greater activation for self-criticism (emotion - neutral) - self-reassurance (emotion - neutral). Peak thresholded at FWE, $p$ <0.05 with K = 110 and coordinates reported in MNI-space. Cluster labelling was used from the AAL toolbox implemented in SPM12.
